# Physiological synchrony promotes cooperative success in real-life interactions

**DOI:** 10.1101/792416

**Authors:** F. Behrens, J. A. Snijdewint, R. G. Moulder, E. Prochazkova, E. E. Sjak-Shie, S. M. Boker, M. E. Kret

**Author notes:** Corresponding Author: M. E. Kret, Leiden University, Institute of Psychology, Cognitive Psychology Unit, Wassenaarseweg 52, Leiden 2333 AK, The Netherlands;.

## Abstract

Cooperation is pivotal for society to flourish and prosper. To ease cooperation, humans express and read emotions and intentions via explicit signals or subtle reflections of arousal visible in the face. Evidence is accumulating that humans synchronize these nonverbal expressions, as well as the physiological mechanisms underlying them, potentially promoting cooperative behavior. The current study is designed to verify the existence of this putative linkage between synchrony and cooperation. To that extent, 152 participants played multiple rounds of the Prisoner’s Dilemma game in a naturalistic dyadic interaction setting. During one round of games they could see each other, and during a second round they could not. The results showed that when people’s heart rate and skin conductance level aligned, they cooperated more successfully. Interestingly, for skin conductance level synchrony to boost cooperation, face to face contact was essential. The effect of heart rate synchrony on cooperation was context independent. Skin conductance level, but not heart rate, tended to closely correlate with changes in pupil size and we discuss how the pupil might provide a window to partners’ sympathetic arousal. These findings show how unconscious mechanisms guide social behavior and emphasize the importance of studying social processes *between* rather than within individuals in real-life interactions.

**Significance Statement:** Human cooperation is remarkable in its scale, complexity, and efficiency. However, whereas people think of themselves as rational agents, they actually partly base their decision to cooperate with someone on gut feelings. These feelings are informed by non-verbal expressions that are picked up implicitly and that synchronize across interaction partners. For the first time, we show that the alignment of people’s arousal over multiple rounds of the Prisoners’ dilemma game predicts cooperative success. Through synchrony, partners converge emotionally, fostering understanding and bonding, which are key ingredients when it comes to successful cooperation. This suggests that successfully cooperating does not depend on individuals, but on the connection *between* individuals, emphasizing the importance of studying social decision-making processes in real-life settings.

## Introduction

Cooperation is one of human society’s core pillars, distinguishing us from other species in its scale and complexity (1–3). However, warfare, genocide and our role in other species’ extinction darken our evolutionary path and characterize a failure to cooperate globally (4). Importantly, world problems or multinational conflicts are not decided on the battlefield, but during negotiations between leaders. When these powerful policy makers are faced with opponents such as during Brexit-negotiations, emotions overrule rationality. How can cooperation be achieved? In order to be able to foster cooperation, we must first understand the mechanisms. The current study takes a step in that direction.

When making decisions, such as whether to cooperate or not, people rely on a variety of nonverbal expressions to communicate and predict others’ intentions (5–7). Cooperation is risky as individuals can take advantage of those investing time and resources, and nonverbal expressions reflecting a person’s benign intents can be determinative for cooperative success. Intriguingly, research has shown that emotional states tend to synchronize between interaction partners on several levels including the behavioral level (8, 9), the neural level (10, 11), and the physiological level (12, 13). Although some of these emotion-induced changes cannot be observed by the naked eye directly, people perceive them indirectly through visual cues such as pupil size, and align their bodily responses accordingly (13). Whether or not physiological synchrony is a mechanism underlying cooperative decisions is a key question that thus far remains unanswered.

Raising awareness of synchronized emotion states has had a vast impact on different disciplines with researchers investigating its clinical (14, 15), developmental (16, 17), evolutionary (18, 19), neural (11, 20), social (21), and cognitive (22, 23) implications. It has been proposed that the *function* of this alignment is to infer the other person’s emotions, to empathize, and to provide subsequent consolation, help, or other prosocial behavior (24). Despite the clear predictions regarding the function of synchrony, studies have thus far only investigated the benefits of synchrony in artificial settings with either participants interacting with virtual characters on a computer screen (25), or two people interacting in cooperative compared to competitive contexts (26–28). Thus far, no research has investigated direct links between synchrony and subsequent cooperative decisions.

Does synchrony lead to cooperation? This pivotal question has never been directly addressed before. By investigating whether cooperative success can be predicted by interpersonal synchrony in a dyadic real-life interaction, we aim to close this knowledge gap. Here, we focus on physiological synchrony because it is implicit, hard to control or regulate, and is a crucial component of emotion processing (29–31). In psychology, the most commonly studied physiological responses are skin conductance level, a purely sympathetic nervous system response, and heart rate, which reflects both sympathetic and parasympathetic nervous system activity (13, 32, 33). Previous research has shown that before people make a decision and, for instance, express that verbally or via a button press in an experiment, that decision is already reflected in their physiology (34, 35). We here focus on these two measures, investigating whether they synchronize between interaction partners and if so, whether that influences the cooperative success of a dyad.

To this end, 152 naïve participants played a modified iterated Prisoner’s Dilemma game in dyads, 30 trials facing each other (allowing for nonverbal communication), and 30 trials with a visual cover between them, constraining them from interacting nonverbally (Panel A in Figure 1). Based on the original Prisoner’s Dilemma game, we developed a novel game where the payoff structure was extended from the classical two options (i.e., to cooperate or to defect) to a 6 × 6 payoff structure. This modification gave us a more fine-grained measure of cooperation (Methods). To quantify physiological synchrony, we conducted a windowed cross-lagged correlation analysis. This method accounts for the non-stationarity of the time series and delays between individuals’ responses reflecting the dynamical nature of the interaction between two participants (Methods). The heart rate analysis included 60 dyads and the skin conductance level analysis 50 dyads which lies in the upper range of sample sizes across studies investigating physiological synchrony (Methods) (32). The aim of the study was twofold: The first aim was to investigate whether physiological synchrony predicts cooperative success. To investigate the robustness of this putative effect, we used two different, widely used physiological measures (i.e., heart rate and skin conductance level). Second, we aimed to confirm that synchrony and its linkage to cooperative success was bound to interactions where partners could see each other.

**Fig. 1.**
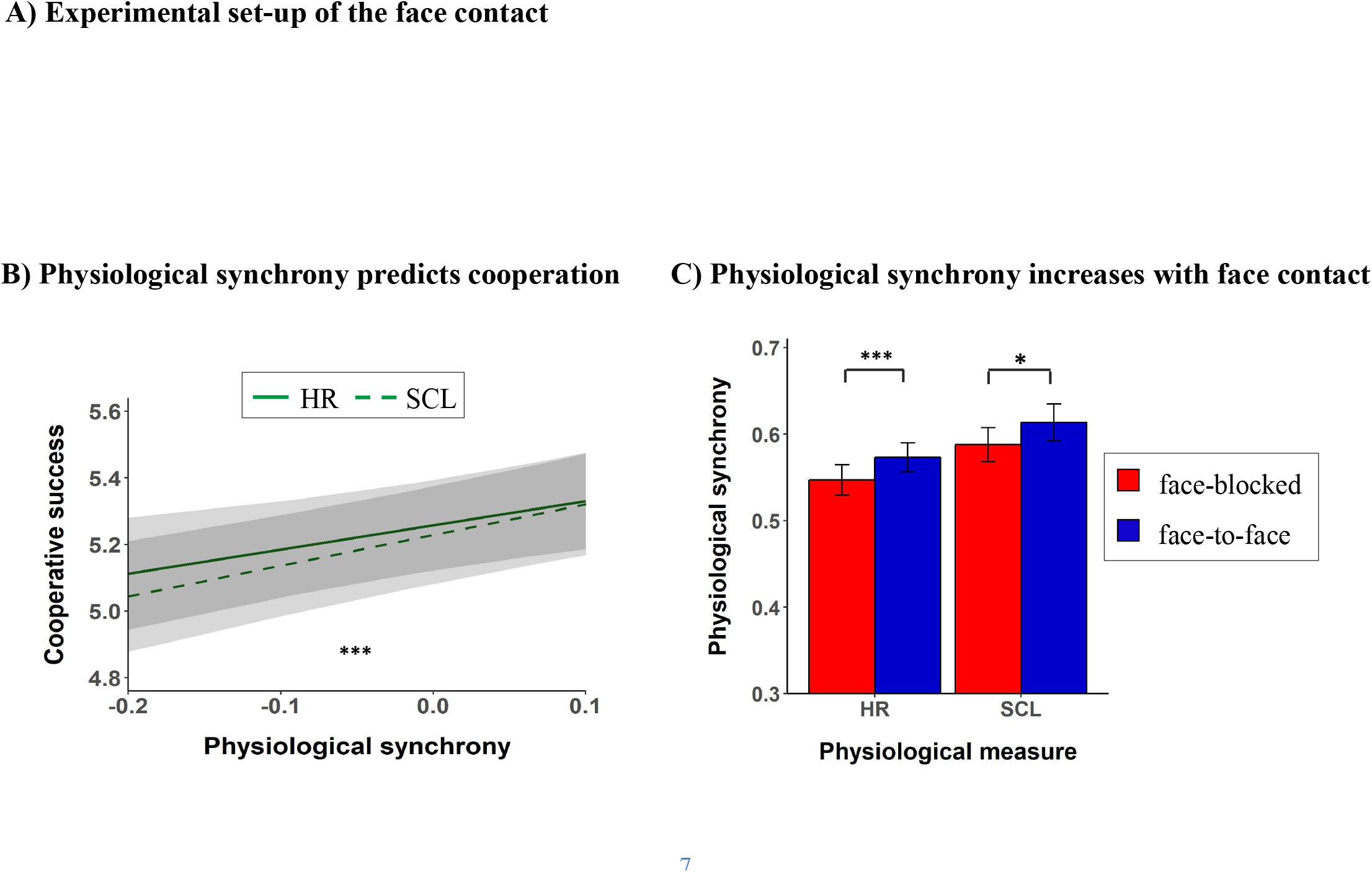
Experimental set-up and results. (A) Dyadic interaction in the face-to-face (left) and face-blocked (right) conditions. (B) Predicted cooperative success based on the synchrony in heart rate (solid) and skin conductance level (dashed). (C) Mean differences between the face-to-face (blue) and face-blocked condition (red) for heart rate and skin conductance level synchrony. The shaded areas in (B) and error bars in (C) represent 95%-confidence intervals. Physiological synchrony is measured by the mean lagged windowed cross-correlation and is grand-mean centered for the analysis displayed in (B) (see Methods for details). Cooperative success is measured by the joint outcome of a dyad per trial in the economic game (range: 4-6 points). HR = Heart rate; SCL = Skin conductance level. * *p* < .05; ** *p* < .01; *** *p* < .001.

## Results

Investigating the joint outcome, the results revealed that stronger synchrony predicted more successful cooperation in both the synchrony in heart rate (*χ*^2^(1) = 8.34, p = .004) and skin conductance level (*χ*^2^(1) = 21.31, *p* < .001; Panel B in Figure 1). Additionally, the effect of synchrony on cooperation was moderated by face contact for skin conductance level synchrony (*χ*^2^(1) = 10.18, *p* = .001). Specifically, the positive relation between skin conductance level synchrony and cooperation was driven by the condition where people could see each other. For the synchrony in heart rate, results yielded no evidence that the effect of synchrony on cooperative success was influenced by face contact (*χ*^2^(1) = 0.21, *p* = .646). Other findings underscore the importance of face contact. Regarding the behavioral responses, participants were more successful in cooperating when they faced each other as compared to when they did not (*χ*^2^(1) = 57.16, *p* < .001; Figure S1) [for similar findings, see (36, 37)]. With respect to physiological synchrony, as predicted, face-to-face contact amplified the level of synchrony in heart rate and skin conductance level (HR: *χ*^2^(1) = 12.87, *p* < .001; SCL: *χ*^2^(1) = 5.57, *p* = .018). See Panel C of Figure 1 for the corresponding plots. Finally, in a control analysis, we compared the level of synchrony from the original dyads with newly generated, randomly matched dyads. Thus, participants were paired with another partner than the one they had actually interacted with in the experiment and we used this analysis to verify that the level of synchrony was due to the interaction rather than the experimental set-up of the study. For both heart rate and skin conductance level, the original dyads showed significantly higher Fisher-Z transformed correlations than the newly generated dyads (HR: *t*(3622.7) = 8.06, *p* < .001; SCL: *t*(3015.5) = 4.38, *p* < .001; see Methods for more details).

## Discussion

For thousands of generations, humans have cooperated with others, familiar or unfamiliar, on unprecedented scales, which has been essential for their survival (1). However, as is clear when opening the newspaper, cooperation also often fails. The core question is: what is the mechanism underlying successful cooperation? The current study gives insight into this question by investigating whether cooperative success can be explained by both partners picking up the nonverbal signals that reflect their physiological arousal, emotionally converging, and consequently fostering mutual understanding and trust. Specifically, the aim of the current study was to investigate the linkage between physiological synchrony and cooperation. For the first time in the literature, we confirmed that physiological synchrony predicts cooperative success in real-life interactions. This link is strengthened when people face each other. Interestingly, this moderation is only evident for skin conductance level synchrony, but not for heart rate synchrony. Furthermore, both physiological synchrony and cooperative success are higher when people face each other, and synchrony levels are higher in real compared to artificially generated dyads. These findings imply that people are able to detect subtle changes in the other person’s face, and react to these changes, which subsequently makes cooperation more successful. Physiological synchrony acts as an unconscious mechanism to guide our behavior and improve the success of close social interactions.

Synchronization is observed on many different levels (13), in young infants (12, 38) and in different species (18, 39). Theoretically, it makes two interaction partners more similar, aligned, and easier to predict which is why they cooperate more efficiently (13, 24). By manipulating a cooperative versus competitive context, previous research showed increased heart rate synchrony (40) and skin conductance synchrony (28) in a cooperative compared to competitive context. The current study builds on this earlier work by showing that, on the basis of synchrony, cooperation could be predicted on a trial-by-trial basis. This new approach better reflects natural situations where multiple small decisions are taken in a row and thus shows the real impact of synchronization on these decisions. Although the current design does not permit strong conclusions regarding causality of the observed effects, it allows cooperation to naturally evolve through genuine interpersonal processes.

Cooperation carries the risk of being exploited by non-cooperators, therefore being able to detect the integrity of another person’s intent is crucial. Although explicit, visible signals such as facial expressions and eye gaze can provide valuable information, these signals are easily faked and do not necessarily reflect a person’s true intentions (5, 41, 42). Physiological responses, on the other hand, are difficult to control and constitute a crucial component of a person’s emotional state and are indicative of social decision-making (7, 31, 43). Synchronizing on the physiological level has been proposed to be able to change the way Person A feels about and behaves towards Person B which is consequently communicated in signals visible to Person A (41). Likewise, if the explicit signals do show the benign intentions, such signals and their mimicry can influence autonomic responses and their synchrony implying a bi-directional interaction between autonomic cues and explicit signals. The influence of visible signals on the synchrony in heart rate and skin conductance level is supported by the current findings that people synchronized to a greater extent when they could face each other compared to no face contact; visible signals could be exchanged in the former but not the latter condition. Thus, we argue that cooperation flourishes when people synchronize their autonomic responses because they align emotional states based on genuine emotional cues that are perceived by interaction partners.

Which visible signals are essential in communicating the autonomic changes in a person is not yet clear. Besides the more pronounced signals such as facial expressions and eye gaze, other subtle, yet visible cues that are closely linked to changes in arousal are pupil dilation and blushing. Both cues have been shown to influence trust, a precursor of cooperation (27, 44). The physiological correlates of blushing are less understood with some studies observing only sympathetic influence and others proposing influences from both components of the autonomic nervous system (45, 46). On the other hand, people have been shown to be sensitive to pupil size changes in another person such that an observer mimics these changes and subsequently increases her trust towards the observed person (27). Interestingly, changes in pupil size have been shown to co-vary directly with changes in skin conductance level, but not heart rate (47). This link is in line with the current findings that for skin conductance level synchrony to boost cooperation, face contact was essential, whereas this was no boundary condition for the heart rate synchrony-cooperation linkage. Future studies should clarify whether pupil dilation might indeed be the channel through which changes within the person are detected and emotional states align between individuals.

The fact that face contact specifically boosts the synchrony-cooperation linkage for the skin conductance level, but not for heart rate is further in line with previous studies linking these measures to behavioral responses. For example, sympathetic synchrony has been shown to elicit perceived similarity between interaction partners (48) and perceived similarity has been shown to foster cooperation (49). Furthermore, the sympathetic changes in skin conductance level have been related to (disadvantageous) decision-making and emotion regulation (7, 34, 50). Given the risk of being exploited during cooperation, one might need increased emotion regulation to control the urge to defect in order to successfully cooperate. “Clicking” with another person on the autonomic level might therefore be an essential component of cooperation. These suggestions are, however, speculative and future research is needed to draw strong conclusions about how different responses are integrated in affecting social decision-making.

The current study has significant implications for studying the intricate dynamics of cooperation. We provide unique evidence that physiological synchrony functions as an underlying mechanism of cooperative success. Studying cooperation in a real-life interaction setting helped us to unfold a new layer of communicative processes that is ignored when investigating social phenomena in computerized, one-person paradigms (51). This new layer incorporates how two bodies communicate on a subtle level that we are not aware of, yet that influences the way we behave towards other individuals. Shedding light onto what makes cooperation successful in healthy interactions can help us understand situations where human interactions fail. Conflict resolution, whether in a conversation, a company or an international collaboration, is dependent on moment-by-moment cooperative tendencies of its individuals. Such tendencies are by virtue reliant on human’s ability to understand each other’s emotions and on the capacity to balance their emotions with one another. Applying this to clinical populations, it has been suggested that the lack of interpersonal exchange of nonverbal signals underlie different deficits evident in autism, social anxiety, and depression which can advance new therapies in these populations (15, 52). Our findings broaden our understanding of the function of synchrony in social behavior and add a hereto forth missing piece to the puzzle of understanding the unconscious mechanisms guiding cooperation.

## Materials and Methods

### Experimental Design

The objective of the study was to investigate whether cooperative success could be predicted based on the physiological synchrony between two individuals in a real-life interaction setting. To this end, two participants played a modified iterated Prisoner’s Dilemma game while their heart rate and skin conductance responses were measured. A mixed-design study was conducted with one within-dyad (Face manipulation) and one between-dyad (Feedback manipulation) variable. In the latter manipulation, people received auditory feedback about their decision or not. However, this manipulation did not influence cooperation (*χ*^2^(1) = 1.29, *p* = .256), and was not the focus of this article. As such, the Feedback manipulation is not discussed and the data were pooled together for all analyses. Regarding the Face manipulation, participants could either see each other’s faces (face-to-face condition) or they could not see each other (face-blocked condition). The dependent variable was cooperation which was measured by means of a modified version of the Prisoner’s Dilemma game (see below). All dyads played 30 rounds of the game in both conditions with the order counterbalanced. During the whole experiment, participants’ heart rate, skin conductance level and eye movements were measured.

### Participants

In total, 152 individuals participated in the study (71 % females, *M*_*age*_ = 23, *SD*_*age*_ = 4.3), who were recruited via the University online recruitment system (SONA) and by approaching people on University ground. A dyad consisted of two same-sex individuals who did not know each other (*N*_*dyads*_ = 76). All participants had normal or corrected-to-normal vision wearing contact lenses (glasses were not compatible with the eye-tracking glasses, see below). They received either course credits or a monetary reward (8€) for participation and could earn an additional maximum of 2€ depending on their performance during the experiment. The study was approved by the Psychology Research Ethics Committee of Leiden University (CEP17-0113/18).

### Missing data

For the behavioral data, three of the 152 participants (=76 dyads) were excluded because they had missing data for 30 or more out of 60 trials. For the physiological data, the decision to exclude data was based on the manual preprocessing of the data. Either the measurement of the physiological responses was erroneous in at least one of the two participants during the whole session or more than 70% of the responses were missing due to local measurement errors in the data. Based on these criteria, 14 dyads had to be excluded. Two additional dyads were excluded because they did not make any eye-contact during the face-to-face condition trials which was verified by means of eye-tracking glasses worn during the experiment. Ten additional dyads were excluded from only the skin conductance level analysis due to measurement errors. Thus, the heart rate analysis included 60 dyads and the skin conductance level analysis 50 dyads which lies in the upper range of sample sizes across studies investigating physiological synchrony (32). In addition, 29 single trials for the heart rate data and three single trials for the skin conductance level data were excluded.

## Materials

### Cooperation game

To measure cooperation, a modified version of the Prisoner’s Dilemma game was used. The general idea of the game is that people can choose between two options (cooperate versus defect) that affect both a person’s own and the partner’s outcome. In particular, if both players cooperate (CC), each player receives more points compared to if both players defect (DD). If one player cooperates and the other defects, the latter receives the highest points possible, while the former receives the lowest points. Hence, the dilemma is to choose between maximizing the own outcome by defecting (which is more advantageous independent of the other player’s choice) or maximizing the joint outcome by cooperating (the highest joint outcome is achieved when both players cooperate). In the current study, the idea of the game stayed the same, but people could choose between six instead of two options (option A-F) creating a cooperation scale (Table 1). For this purpose, we built two boards where participants could put a pawn on the response matrix to indicate their response. That response incorporated two choices: (1) the level of willingness to cooperate; moving from the left (option A) to the right (option F) on the x-axis, the willingness to cooperate increased with option A reflecting complete defection and option F reflecting complete cooperation; (2) what the participant thought the other person would choose on that trial; moving from the bottom (option A) to the top (option F) on the y-axis indicates that the participant expected the partner to cooperate more. Hence, the highlighted options in the four corners in Table 1 reflect the payoff structure of a traditional Prisoner’s Dilemma game, but the extended matrix shows the innovative structure designed for the current experiment.

**Table 1.** Payoff structure of the current study (bold numbers were not highlighted during the experiment).

|  |  |  |  |  |  |  |  |
| --- | --- | --- | --- | --- | --- | --- | --- |
| *Other* | F | **4.0 - 1.0** | 3.8 - 1.4 | 3.6 - 1.8 | 3.4 - 2.2 | 3.2 - 2.6 | **3.0 - 3.0** |
|  | E | 3.6 - 1.2 | 3.4 - 1.6 | 3.2 - 2.0 | 3.0 - 2.4 | 2.8 - 2.8 | 2.6 - 3.2 |
|  | D | 3.2 - 1.4 | 3.0 - 1.8 | 2.8 - 2.2 | 2.6 - 2.6 | 2.4 - 3.0 | 2.2 - 3.4 |
|  | C | 2.8 - 1.6 | 2.6 - 2.0 | 2.4 - 2.4 | 2.2 - 2.8 | 2.0 - 3.2 | 1.8 - 3.6 |
|  | B | 2.4 - 1.8 | 2.2 - 2.2 | 2.0 - 2.6 | 1.8 - 3.0 | 1.6 - 3.4 | 1.4 - 3.8 |
|  | A | **2.0 - 2.0** | 1.8 - 2.4 | 1.6 - 2.8 | 1.4 - 3.2 | 1.2 - 3.6 | **1.0 - 4.0** |
|  |  | A | B | C | D | E | F |
|  |  | *You* |  |  |  |  |  |

*Note. The first number refers to the points earned by “You”.*

### Physiological data acquisition and preparation

Throughout the experiment, four physiological responses were measured on both participants: heart rate (HR), skin conductance level (SCL), zygomaticus major (smiling muscle) and eye movements by means of electrocardiography (ECG), electrodermal activity (EDA), electromyography (EMG), and eye tracking glasses, respectively. The former three were recorded wirelessly with the MP150 BIOPAC data acquisition system and sampled at 2000 Hz. The EMG data contained many artifacts where the source could not be identified and the shape of the artifacts did not allow for clear distinction between artifacts and responses. Therefore, the facial expression data were not included in this paper.

For the analyses, the preprocessed heart rate and skin conductance level measures were down-sampled to 20 Hz. The software AcqKnowledge (AcqKnowledge v. 4.4; BIOPAC Systems Inc.) was used to record and sync the signals from the physiological signals, the event markers from E-Prime which was used to present the instructions and lock the behavioral responses, and markers sent by the eye tracking glasses.

#### Heart rate

To measure participants’ heart rate, electrodes were attached on the left and right side of the abdomen and on thorax below the right collar bone. To process the data, an in-house developed software, PhysioData Toolbox (53), was used offline. The signals were band-filtered with a cut-off of 1 Hz and 50 Hz. The R-peaks that were automatically detected by the software were afterwards visually inspected and manually corrected in case of missed or incorrect R-peaks. To still generate a smooth and continuous heart rate signal, interbeat intervals (IBI) were linearly interpolated in these locations. Participants with less than 30% coverage of the sum of the IBIs relative to the duration of the time signal were excluded. The signal used for the analyses was heart rate which was measured in beats-per-minutes.

#### Skin conductance level

Two electrodes were attached on the intermediate phalanges of the index and ring finger of the non-dominant hand. To improve the quality of the signal, there was a time interval of around 15 minutes between the attachment of the electrodes and the beginning of the data collection. The skin conductance level measures were low-pass filtered with a cut-off of 5 Hz and subsequently visually inspected for artifacts using the PhysioData Toolbox (53).

#### Eye movements

Participants were wearing Tobii Pro Glasses 2 to track their eye movement and to verify whether they were looking at each other during the face-to-face condition trials. Fixation points were manually coded in Tobii Lab Pro (version 1.64, 2017). Trials in which participants were not at least once looking at the face of the other person were excluded.

### Procedure

Before participants came to the lab, they received information about the study and filled out three questionnaire about empathy [Interpersonal Relation Index; IRI; (54)], social anxiety [Liebowitz Social Anxiety Scale; LSAS; (55)], and social value orientation [SVO; (56)]. Upon arrival at the lab, participants signed an informed consent in separate rooms and a female researcher attached the electrodes for measuring heart rate, skin conductance level, and facial expressions (Methods). Next, participants filled out the Positive And Negative Affect Scale [PANAS; (57)] and read the instructions for the social dilemma game. Their understanding of the game was checked with multiple choice questions which were discussed in more detailed when answered incorrectly. Afterwards, both participants sat on a table in front of each other with a wooden board between them such that they could only see each other’s faces. Finally, the eye tracking glasses were calibrated, the researcher left the room and started the experiment.

After three practice trials (face-to-face condition), participants played the game two times 30 rounds in the face-to-face and face-blocked condition. The order of starting in one or the other condition was counterbalanced. To block nonverbal communication in the latter condition, a visual cover was placed on top of the wooden board. The sequence of the trial was as follows with auditory instructions given via speakers: First, participants were instructed to look at each other (look at the cross in front of them [drawn on the visual cover] in the face-blocked condition). After four seconds, they were asked to look down and make a decision. When both individuals made their decision, they either heard that they have both made a decision (no feedback condition) or heard how many points each player received based on their choices (feedback condition). As mentioned above, the role of feedback is not discussed here.

After each session, participants filled out a visual analogue scale (VAS) about their current feelings and experiences. After the second session, participants were separated again in different rooms where they filled out the Desire for Future Interaction scale [DFI; (58)], read the debriefing form. Finally, they were paid and thanked for participation.

### Statistical Analysis

During the study, different questionnaires about the participants’ characteristics and current mood and experiences were measured as mentioned in the Procedure. These data were not the focus of the current article and are not discussed any further.

### Behavioral data

We hypothesized that face contact would increase the joint outcome, i.e. cooperative success. Specifically, cooperative success was measured as the points both players earned together which ranged from 4.0 to 6.0 points. The Face condition variable was coded 0 = face-blocked condition and 1 = face-to-face condition. We conducted a multilevel linear regression analysis with dyads added as a random intercept effect. The inclusion of the random effect was verified by running an empty model consisting of the random effect only and calculating the intra-class correlation which quantifies how much dependency there is in the data. Significance of fixed effects was determined by means of model fit comparisons between the model including and excluding the effect of interest. The Likelihood Ratio Test (LRT) was used as the reference. It calculates the difference in deviance between the full and reduced model and follows a *χ*^2^-distribution with the difference in number of estimated parameters between models as the degrees of freedom. The significance level of .05 was applied. Dyads with more than 50% missing data (more than 30 trials) were excluded.

### Physiological data

We conducted a lagged windowed cross correlation analysis (59) to quantify physiological synchrony for the heart rate and skin conductance level measures separately. Based on this analysis, we obtained a measure of the strength of synchrony for each Face condition per dyad.

#### Quantifying physiological synchrony

Two methods that take non-stationarity into account are lagged windowed cross-correlation (59) and recurrence quantification analysis (60). The latter method is frequently used which has the advantage of having very few assumptions. However, the disadvantage is that it determines synchrony on a binary scale of moments being classified as either synchronized or not. The former method, albeit constraint by more assumptions, has the advantage of differentiating the degree of synchronization by quantifying it on a continuous (correlation) scale. Additionally, we feel that windowed cross-correlation is more intuitive to interpret. Consequently, we decided to apply this method which provides measures of the strength of synchrony and its variability.

The objective of the lagged windows-cross correlations analysis (59) is to calculate the strength of association between two time series while taking into account the non-stationarity of the signals and the lag between responses, that is, to consider the dynamics of a dyadic interaction. Specifically, the time series are segmented into smaller intervals, calculating the cross-correlation for each segment. This allows the means and variances to differ between segments accounting for non-stationarity. This is important as the level of synchrony may change during the experiment, sometimes having moments of strong synchronization while during other times responding less strong to one another. Additionally, as the strength of association between two time points may differ depending on how far apart they are from each other, the segments are moved along the time series by an increment such that two adjacent segments overlap. Hence, segmenting the time series into smaller intervals and partially overlapping these intervals while moving along the time series provides a better estimate of the local strength of association between the physiological signals of two participants.

Besides the dynamics in the strength of synchronization during the course of the experiment, participants differ in how fast one might respond to a certain event or the other person. In other words, participants might not always be perfectly “in sync” whereby one participant might sometimes respond to the other person or vice versa introducing a delay between the responses of two individuals. To account for this, for each segment, the signals of the two participants are lagged in relation to one another. Specifically, the signal of participant 1 is kept constant while the signal of participant 2 is shifted more and more by a specified lag increment until a maximum lag is reached. Next, the same procedure is performed the other way around with participant 2 being kept constant. The maximum lag determines what is still considered synchrony. For example, if the maximum lag is four seconds, responses from two participants that are four seconds apart from each other are still considered synchronized. On the other hand, if one participant reacts to a certain event and the other participant shows a response 5 seconds later, it is not considered a response to the same event anymore and therefore does not count as synchrony. Based on this approach, there are four parameters that need to be determined: (1) the length of each segment, referred to the window size *w*_*max*_; (2) the increment with which the segments are moved along the time series, the window increment *w*_*inc*_; (3) the maximum with which two segments can be lagged from one another, the maximum lag *τ*_*max*_; and (4) the increment with which two segments are lagged from each other, the lag increment *τ*_*inc*_. We determined the parameters following an extensive process by comparing previous studies using similar statistical methods, by looking at what is physiologically plausible given the time course of the physiological signals and by employing a data-driven bottom-up approach where we investigated how changing the parameters affected the outcomes using a different dataset. As expected, the absolute values of the synchrony measures varied depending on the parameters, but as supported by (61), the relative results were not affected (e.g. a dyadic manifesting relatively high synchrony showed such tendency for the different parameters). Based on these three factors, we set the parameters as follows: the window size was 8 seconds (160 samples), the window increment was 2 seconds (40 samples), the maximum lag was 4 seconds (80 samples) and the lag increment was 100ms (2 samples).

Calculating the cross correlations of each lag for each window segment generates a result matrix with each row representing one window segment and each column indicating a lag. The middle column represents the cross-correlation with a lag of zero, while the first and last column contain the cross-correlations for the maximum lag of participant 1 and 2. Hence, the number of columns in the result matrix is (2* *τ*_*max*_ / *τ*_*inc*_) + 1. The number of rows is given by (*N* − *w*_*max*_ − *τ*_*max*_)/ *w*_*inc*_, with N being the number of observations in the whole time series.

Based on this result matrix, a so-called peak picking algorithm is applied. For each segment (i.e., each row in the matrix), the maximum cross-correlation across the lags is detected closest to the zero-lag (i.e., across all columns in a given row). If that maximum correlation is preceded and followed by smaller correlations, it is marked as a peak. For example, if participant 2 synchronizes with participant 1 with a lag of one second, the cross-correlations will become higher the closer the segments from the two participants are shifted towards the point where they are one second apart from each other. When the two signals are lagged by exactly one second the cross-correlation is highest (the peak). If the signals are lagged further away from each other, the cross-correlation decreases again. If, however, a peak cannot be detected, the algorithm assigns a missing value for that segment. This might be the case, for example, if people do not respond to an event or to each other (e.g., both participants wait and do nothing). The peak picking algorithm outputs a matrix with two columns, containing the value of the maximum cross-correlation (the peak) and the corresponding lag at which the peak cross-correlation is detected. The output has the same number of rows as the result matrix as it searches for a peak cross-correlation for each window segment.

Both the windowed cross-correlations and the peak picking algorithm are conducted four times per dyad, once for the heart rate responses and once for the skin conductance level responses for the face-to-face session and for the face-blocked condition resulting in *N*_*dyads*_ * 4 result and peak picking matrices. Finally, the mean of the peak cross-correlations of all window segments (i.e., all rows of the peak picking matrix) is calculated for both physiological measures per Face condition per dyad as the measure of synchrony and is grand-mean centered for the analysis predicting cooperative success.

#### Hypothesis testing

Based on the synchrony measures we conducted two analyses per physiological response to (i) investigate whether synchrony is influenced by the face contact manipulations, and (ii) test whether the joint outcome can be predicted based on synchrony and on whether people could see each other or not. For both analyses, multilevel linear regression analyses were performed with the same procedure as for the behavioral data. Regarding the first part, Face condition was added as the predictor and the synchrony measure for heart rate and skin conductance level responses as the outcome variables. For the second part, we included the synchrony measures as the predictor and the cooperative success as the outcome variable. In a last step, we added the two-way interactions between Face condition and the synchrony measure.

#### Control analysis

Because the heart rate and skin conductance level will always show a certain level of synchrony between participants due to the nature of the signals and the experimental set-up, we conducted a control analysis to show that synchrony was elevated due to the interaction itself. Specifically, we compared the original dyads with newly generated dyads (Player 1 from Dyad_i_ and Player 2 from Dyad_i+1_). Because the trial length varied (there was no time restriction for making a decision), each trial was cut to the shorter trial of the newly generated dyad. Subsequently, the correlation for the heart rate and skin conductance was calculated per trial and dyad. Finally, we ran an independent t-test on the Fisher-Z-transformed correlation values between the original and the newly generated dyads.

## Acknowledgments

We thank the students who have assisted in collecting the data for this study.

## Funding

This research was supported by the Leiden University Fund / Mr.J.J. van Walsem Fonds W18205-5-95 (to F.B.), Netherlands Science Foundation 016.VIDI.185.036 (to M.E.K.), and the Talent Grant 406-15-026 from Nederlandse Organisatie voor Wetenschappelijk Onderzoek (to M.E.K. and E.P.);

## Author contributions

Conceptualization: F.B., M.E.K., J.A.S, E.P.; Data curation: F.B., J.A.S; Formal analysis: F.B., R.G.M., S.M.B; Investigation: F.B., J.A.S; Methodology: F.B., R.G.M, S.M.B, E.P, Software: E.E.S.S., Supervision: M.E.K, S.M.B; Visualization: F.B.; Writing – original draft: F.B.; Writing – review & editing: J.A.S., R.G.M., E.P., E.E.S.S., M.E.K.;

## Competing interests

Authors declare no competing interests.

## Data and materials availability

All data, code, and materials that are associated with this paper and used to conduct the analyses will be uploaded and made accessible on the Leiden University archiving platform DataverseNL when published.

## Supplementary Materials

Figure S1

## Auxiliary Supplementary Materials

Data file [DataFile_FBehrens_et_al.csv]

R script with analyses performed in this manuscript [Analyses_FBehrens_et_al.r]

## Notes

#### Summary of Updates

[Preprint] added to title for clarity

