## Supplementary Materials - Figure S1 for "Physiological synchrony promotes cooperative success in real-life interactions"

**Fig. S1.**

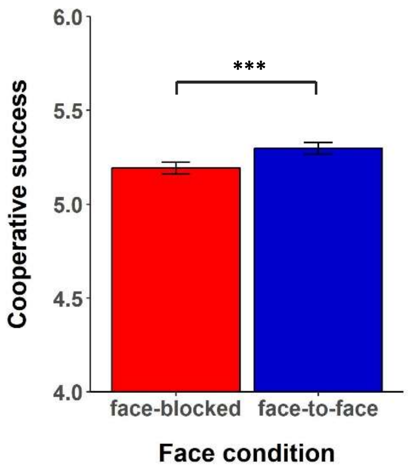


**Fig. S1.** The cooperation success rate for the face-blocked and face-to-face conditions with error bars representing 95%-confidence intervals. * $p < .05$; ** $p < .01$; *** $p < .001$.
